# Parkinson’s Disease-vulnerable and -resilient dopamine neurons display opposite responses to excitatory input

**DOI:** 10.1101/2025.06.03.657460

**Authors:** Lotfi C. Hadjas, Grace J. Kollman, Lexe Linderhof, Mingxuan Xia, Syon Mansur, Martine Saint-Pierre, Byung Kook Lim, Edward B. Lee, Francesca Cicchetti, Rajeshwar Awatramani, Nick G. Hollon, Thomas S. Hnasko

## Abstract

Dopamine (DA) neurons of the substantia nigra (SN) are essential for motor control and selectively degenerate in Parkinson’s disease (PD). However, DA neurons are molecularly heterogeneous, with some showing greater vulnerability and others resilience. Here, we show that the DA subtype marker *Anxa1*, identified in mice, labels PD-vulnerable DA neurons in human SN. Using mice, we found that excitatory inputs from subthalamic (STN) and pedunculopontine (PPN) nuclei evoked frequency-dependent excitation in SN GABA neurons, but complex multiphasic DA neuron responses, suggesting heterogeneous DA subtype responses. Indeed, excitatory inputs evoked differential DA responses in striatal subregions, an increase in caudal striatum, but inhibition followed by rebound in dorsolateral striatum. Additionally, PD-resilient Vglut2+ DA neurons were excited by STN/PPN input, while vulnerable Anxa1+ DA neurons were inhibited. These findings demonstrate that DA subtypes are embedded in distinct functional networks, suggesting that some therapeutic interventions may differentially impact vulnerable and resilient DA subtypes.

## Introduction

Parkinson’s Disease (PD) is a neurodegenerative disorder characterized by cardinal motor symptoms including resting tremor, bradykinesia, and rigidity. These clinical features are primarily caused by the progressive loss of dopamine (DA) neurons in the substantia nigra pars compacta (SNc) (Surmeier et al. 2017; Hornykiewicz 2002). However, not all DA neurons are equally vulnerable. Neuropathological studies have indicated that DA neurons located in ventral SNc, particularly those containing neuromelanin, are disproportionately affected in PD, suggesting heterogeneity within the SNc DA population. This concept has been furthered by work showing that some genes and proteins are differentially expressed in surviving SNc DA neurons in PD compared to non-PD cases (Brichta & Greengard 2014; Giguère et al. 2018; Kamath et al. 2022). For example, DA neurons expressing *CALB1* or *SLC17A6* tend to be overrepresented in PD, suggesting reduced vulnerability (Yamada et al. 1990; German et al. 1992; Damier et al. 1999; Steinkellner et al. 2022); whereas DA neurons expressing *SOX6* or *ALDH1A1* tend to be underrepresented implying increased PD-vulnerability (Liu et al. 2014; Pereira Luppi et al. 2021).

These molecular markers are also expressed in rodents, where they are found in distinct parts of the SNc and can be used as molecular signatures of DA neuron subtypes (Poulin et al. 2014; La Manno et al. 2016; Hook et al. 2018; Gaertner et al. 2022; Yaghmaeian Salmani et al. 2024). These subtypes are associated with distinct anatomical and functional properties. For example, genes associated with DA neuron resilience including *Calb1* or *Scl17a6* (the gene encoding the vesicular glutamate transporter, VGLUT2) which are expressed in overlapping populations of DA neurons concentrated in dorsal and lateral SNc that densely innervate the dorsomedial and tail of striatum (TS) (Menegas et al. 2015; Menegas et al. 2018; Poulin et al. 2018). In contrast, PD-vulnerable *Aldh1a1*-expressing DA neurons are most abundant in ventral SNc and project to dorsal striatum (Poulin et al. 2018; Wu et al. 2019) Recent mouse studies have identified *Anxa1* as subset of *Aldh1a1* DA neurons occupying the ventral-most SNc that project densely to dorsal striatum (DS), and as a maker of vulnerable SNc DA neurons in PD-mouse models (Azcorra et al. 2023; Fushiki et al. 2024; Mantas et al. 2024). In addition to these anatomical differences, resilient and vulnerable SNc DA subtypes have strikingly different patterns of activity associated with spontaneous locomotion. Together, these findings highlight the existence of distinct SNc DA subtypes with differential PD-vulnerability, molecular identities, striatal targets, and in vivo activity patterns. This raises the question of how distinct in vivo activity patterns can be generated in DA subytpes.

Among the excitatory afferents to the SN, the subthalamic nucleus (STN) and pedunculopontine nucleus (PPN) are notable due to their established roles in motor control and their importance to PD. The STN is an important target for deep brain stimulation (DBS) in PD patients and sends dense excitatory projections to the GABAergic substantia nigra reticulata (SNr), as well as to DA neurons in SNc (Bevan 2016; Prasad & Wallén-Mackenzie 2024). PPN is defined by its cholinergic neurons but is highly heterogeneous and contains abundant glutamatergic neurons implicated in motor control, including an ascending projection to both SNr and SNc (Mena-Segovia & Bolam 2017; Leiras et al. 2022). However, whether excitatory STN and PPN inputs differentially engage SNc DA neuron subtypes in vivo remains unknown.

In this study, we first sought to test whether the DA subtype marker *Anxa1*, identified in the ventral SNc of mice, is present in human SNc. We found that indeed *ANXA1* is expressed in human DA neurons and is dramatically de-enriched in PD, suggesting it is a marker of highly vulnerable DA neurons. We next aimed to understand how PD-resilient Vglut2+ and PD-vulnerable Anxa1+ DA neurons are controlled by excitatory inputs in mice. We used optogenetics to stimulate either STN or PPN glutamatergic inputs while making in vivo measurements of either DA release across striatal sites, or calcium activity in SNc DA subtypes. We found that activating these glutamatergic inputs increased calcium activity of PD-resilient Vglut2+ DA neurons and increased DA release in TS. Conversely, activating these excitatory inputs reduced activity in Anxa1+ DA neurons and reduced DA release in DLS. These inhibitions were followed by apparent rebound activation, consistent with a mechanism of feed-forward inhibition. These data indicate that DA subtypes are embedded in distinct neural circuits and suggest new mechanisms by which PD-resilient and PD-vulnerable DA subtypes can be differentially controlled by excitatory afferents.

## Results

### *ANXA1* is expressed in human SNc and marks a PD-vulnerable DA subtype

To determine whether *ANXA1* is a marker of a DA neuron population vulnerable in PD, we performed a duplex chromogenic RNAscope assay on SNc samples from PD cases and age-matched non-PD controls (**Table S1**), labeling for *ANXA1* plus the DA marker *TH* (**Figure 1A-C**). In parallel, we labeled sections from the same cases for *TH* plus either *CALB1* or *ALDH1A1*, allowing us to make comparisons between these DA subtype markers (**Figure 1B, S1A-F, Table S2**). We also cross-validated our approach using immunofluorescence to detect TH/ALDH1A1, including in a subset of the cases that were assessed by RNAscope (**Figure S1G-I, Table S3**).

**Figure 1.**
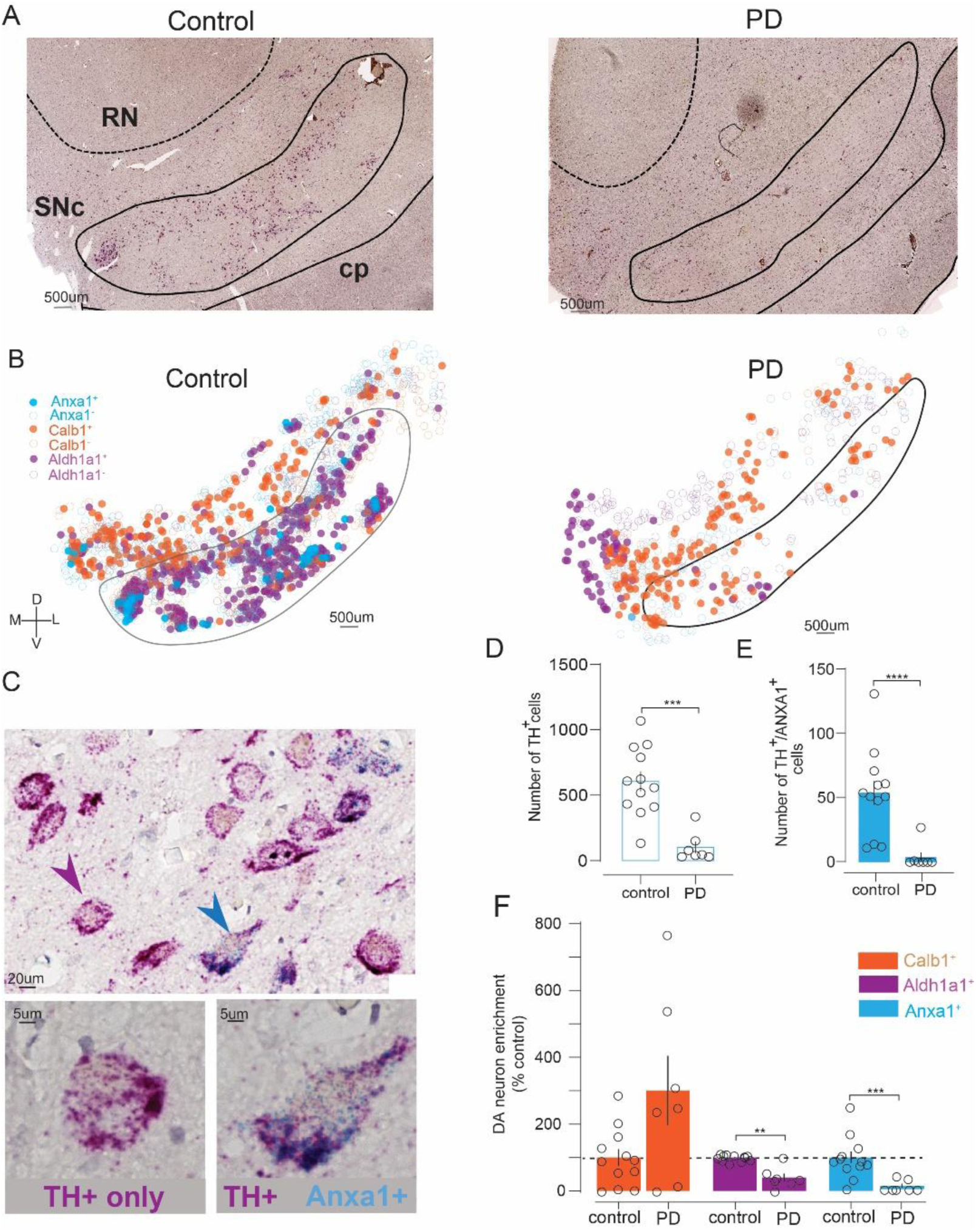
*ANXA1* is expressed in human SNc and marks PD-vulnerable DA neurons. **A)** Representative images showing *TH* (magenta) and *ANXA1* (blue puncta) mRNA in SNc from PD and non-PD control cases. Abbreviations: SNc, substantia nigra pars compacta; RN, red nucleus; cp, cerebral peduncle. **B)** Distribution of *ANXA1*⁺ (blue), *CALB1*⁺ (orange), and *ALDH1A1*⁺ (purple) DA neurons within demarked SNc and surrounding area of the same example control or PD cases. Axes: D, dorsal; V, ventral; M, medial; L, lateral; scale: 500 μm. **C)** Zoomed images showing dual *ANXA1/TH* (blue arrowhead) and *TH*-only neuron (magenta arrowhead) in control SNc. **D)** *TH*⁺ neuron counts were reduced in PD (n=7) versus controls (n=12). **E)** *TH*⁺/*ANXA1*⁺ neuron counts were also reduced in PD. **F)** Relative to controls, *CALB1*⁺ DA neurons were preserved in PD, while *ALDH1A1*⁺ and *ANXA1*⁺ DA neurons were de-enriched in PD. Mann-Whitney or unpaired t-test, **p<0.01, ***p<0.001, ****p<0.0001.

*ANXA1* was identified by blue/green precipitates and was found in a subset of *TH+* (magenta) SNc neurons. Rarely, if ever, was *ANXA1* identified in SNc neurons devoid of *TH* signal. This indicates that *ANXA1* is a DA subtype marker in human SNc. Distribution maps generated for each case revealed that *ANXA1* expression was concentrated in ventral regions of the SNc, especially compared to *CALB1*+ DA neurons (**Figure 1B**). As expected, the total number of *TH*-expressing neurons was significantly decreased in PD cases compared to controls (Mann-Whitney’s test, U=2.0, p=0.002) (**Figure 1D**). The number of *ANXA1+* DA neurons was also significantly reduced in PD cases (U=1.0, p<0.0001) (**Figure 1E**). Consistent with prior reports (Yamada *et al*., 1990; Pereira Luppi *et al*., 2021; Del Rey *et al*., 2024), the number of *CALB1+* DA neurons was not significantly affected, though a trend was observed (U=19.5, p=0.08) (**Figure S1B-C**), while *ALDH1A1*+ DA neurons were significantly reduced in PD (U=1.0, p<0.0001) (**Figure S1E-F**). Moreover, when we assessed the fraction of remaining DA neurons expressing each of the subtype markers in PD relative to control (**Figure 1F**) we found stark differences, with a trend toward enrichment of *CALB1*+ DA neurons (301 +/− 103 %control, unpaired t-test, t=1.9, p=0.10) compared to the significant de-enrichment observed for *ALDHA1A1+ DA neurons* (41 +/− 12 %control, t=4.9, p=0.002), and especially *ANXA1+* DA neurons which were entirely missing in most PD cases (10 +/− 7 %control, Mann-Whitney, U=3.0, p=0.0003). These results indicate that *ANXA1* is expressed in human SNc DA neurons and serves as a new marker of a DA population that is highly vulnerable in PD.

### STN and PPN activation increase SNr GABA neuron activity in vivo

We next aimed to understand how SN cell types respond to excitatory input from STN and PPN in vivo, using mice. We first tested how these excitatory projections alter activity in SNr GABA neurons, because SNr is known to be a major output of both STN and PPN, and SNr GABAergic neuronal activity can inhibit SNc DA neurons (Ammari et al. 2010; Ferreira-Pinto et al. 2021; Ji et al. 2023; Lin et al. 2024).

To activate excitatory inputs from STN or PPN to SNr while simultaneously recording SNr GABA neuronal activity, we used VGLUT2-Cre crossed with VGAT-Flp (vesicular GABA transporter) mice. Adeno-associated virus (AAV) injections achieved Cre-dependent expression of ChrimsonR in either STN or PPN, and Flp-dependent expression of the calcium indicator GCaMP6f in SNr GABA neurons. An optical fiber was implanted in SNr to enable both stimulation of STN or PPN inputs and recording from SNr GABA neurons (**Figure 2A, D**). We applied stimulation trains ranging from 5 to 40 Hz sustained for three different durations (0.25, 1, or 4 s). Post-hoc assessments were made in all mice to validate AAV-mediated expression and optical fiber placements (**Figure S2**).

**Figure 2.**
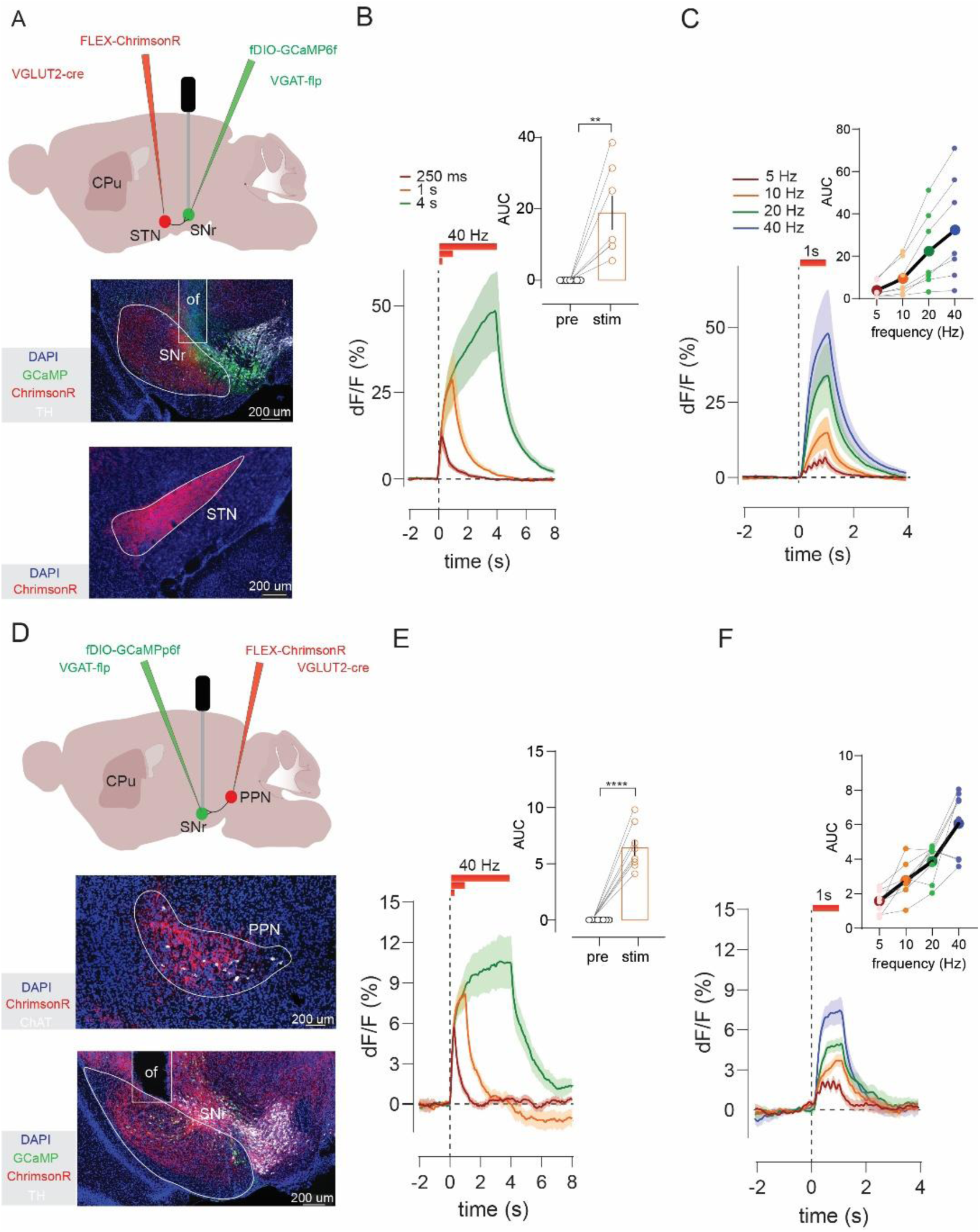
STN and PPN activation increases SNr GABA neuron activity *in vivo*. **A)** Experimental strategy for *in vivo* stimulation of STN glutamate projections to SN and record evoked calcium activity in SNr GABA neurons. Histology shows ChrimsonR-tdTomato in STN and GCaMP plus optic fiber (of) placement in SNr. **B)** Optogenetic stimulation of STN inputs (n=7) at 40 Hz for varying duration increased GABA neuron activity; compared for the 1s train in inset. **C)** One-second STN stimulation (n=7) at increasing frequencies (5, 10, 20, and 40 Hz) produced frequency-dependent increases in GABA neuron responses; compared in inset. **D**) Same as **A)** but for PPN inputs. **E)** Same as B), but for PPN stimulation (n=8). **F)** Same as **C)** but for PPN stimulation (n=8). Paired t-test or Wilcoxon, **p<0.01, ***p<0.001, ****p<0.0001.

Optogenetic stimulation of either STN or PPN led to a significant increase in SNr GABA neuron activity (paired t-test, STN: t=3.8, p=0.008; PPN: t=9.2, p<0.0001) that persisted for the duration of the stimulus (**Figure 2B, E**). Furthermore, the amplitude of the response scaled with the frequency, increasing from 5 to 40 Hz (Friedman’s test, STN: F=21.0, p=0.0001; PPN: F=18.2, p=0.0004) (**Figure 2C, F**). These results confirm that glutamatergic inputs from STN and PPN increase the activity of SNr GABA neurons and provide a basic validation of our approach. We next investigated the effect of STN or PPN stimulation on activity of SNc DA neurons.

### STN and PPN activation drive multiphasic responses in DA neurons

To stimulate STN or PPN inputs and record the activity of the global population of SNc DA neurons we used VGLUT2-Cre to express ChrimsonR in either STN or PPN, with DAT-Flp (dopamine transporter) to express GCaMP6f in SNc DA neurons, and an optic fiber implant targeting SNc (**Figure 3A, E**).

**Figure 3.**
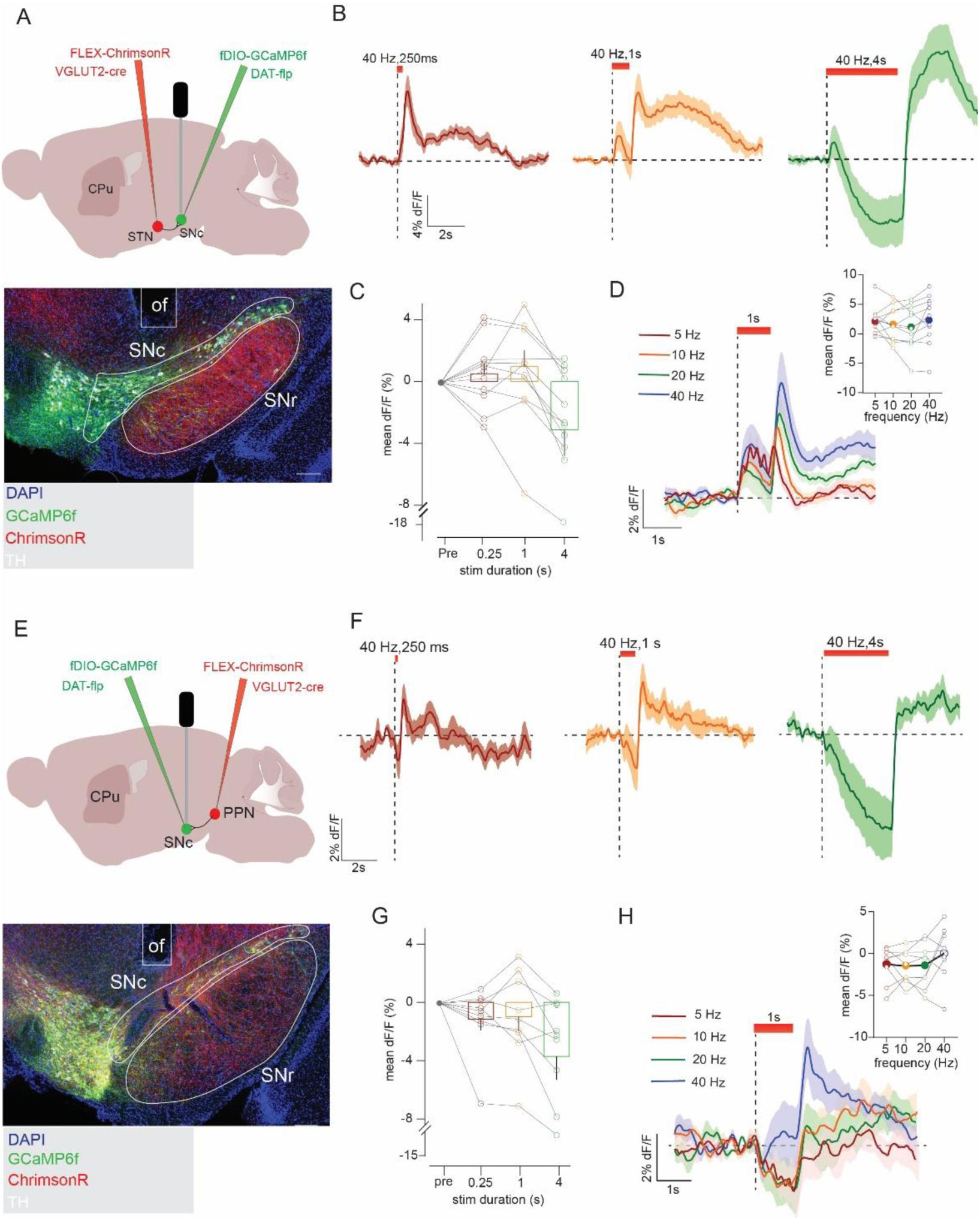
STN and PPN activation drive variable multiphasic pan-DA neuron responses. **A)** Experimental strategy to stimulate STN glutamate projections to SN and record calcium activity in SNc DA neurons. Histology shows ChrimsonR-tdTomato in STN plus GCaMP and optic fiber (of) in SNc**. B)** Optogenetic stimulation of STN (n=11) inputs at 40 Hz for varying duration evoked variable, multiphasic responses in SNc DA neurons. **C)** Quantification of responses (mean ΔF/F during stim train) revealed increases or decreases in DA activity that varied across train duration, **D**) but with no significant (ns) differences as a function of frequency. **E**) Same as A) but for stimulation of PPN projections. **F**) Same as B) but with PPN (n=9). **G,H**) Same as **C,D)** but for PPN.

Stimulation of STN produced multiphasic responses that varied considerably across both the duration and frequency of stimulation (**Figure 3B-D**). For example, a brief (250 ms) 40 Hz train of STN inputs elicited a net increase in DA neuron activity that peaked after stimulus offset (**Figure 3B**; latency to peak, 0.50 +/− 0.03 s). When the train was extended to 1 or 4 s, we observed an initial increase in activity followed by inhibition (Friedman’s test, F=13.0, p=0.005), plus an apparent rebound increase in DA neuron activity at stimulus offset. Notably, the amplitude of the responses evoked during stimulation did not scale neatly with frequency (Friedman’s test, F=4.2, p=0.24) (**Figure 3D**).

Stimulation of PPN inputs led to similarly variable and dynamic DA responses, though they were less likely to provoke the transient activation observed with STN stimulation (Varying durations: Friedman’s test, F=11.5, p=0.009; Varying frequencies: RM one-way ANOVA, F_2,15_=1.35, p=0.28) (**Figures 3F-H**). The complex responses evoked by STN or PPN stimulation in DA neurons are in striking contrast to the sustained, frequency-dependent increases observed in SNr GABA neurons (**Figure 2**). We hypothesized that these complex and variable DA responses were a consequence of the intrinsic heterogeneity within the global DA population. We therefore began testing how discrete DA circuits and subtypes behaved in response to the same patterns of stimulation.

### STN and PPN inputs evoke opposite patterns of DA release in striatal subregions

To investigate DA subtype responses, we first leveraged the known projection topography of DA cell types. PD-resilient Vglut2+ DA subtypes project most densely to TS, whereas PD-vulnerable Anxa1+ DA neurons heavily innervate DLS (Menegas et al. 2015; Poulin et al. 2018; Azcorra et al. 2023). To investigate the effects of STN and PPN inputs on DA release in these regions, we used VGLUT2-Cre mice to express ChrimsonR in STN or PPN and implanted an optic fiber in SNc, as in the experiments described above. We also expressed the genetically encoded DA sensor GRAB_DA2m_ in DLS or TS where we implanted a second fiber. This approach allowed us to stimulate STN or PPN terminals and record DA release in DLS or TS (**Figure 4A-B**).

**Figure 4.**
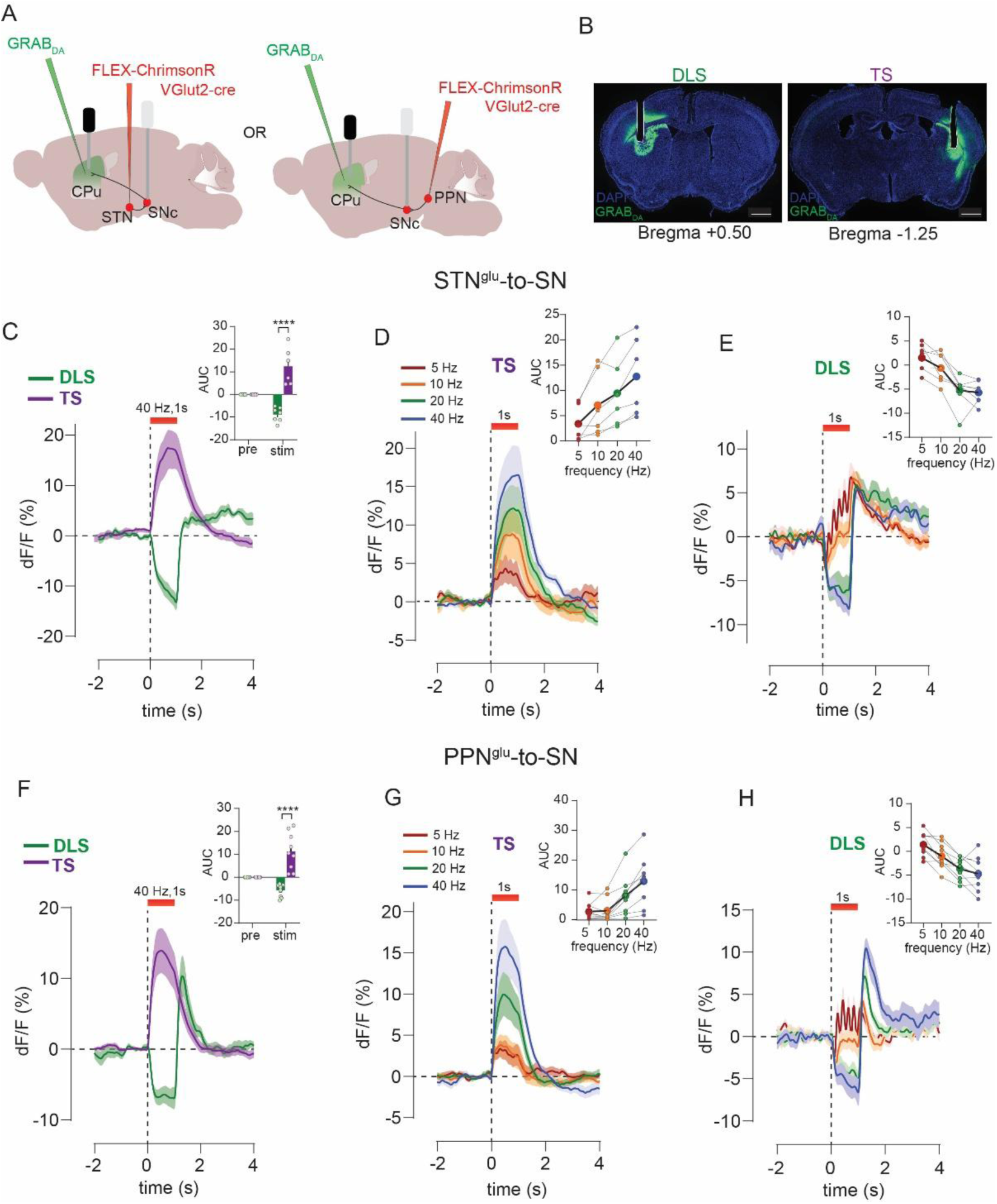
STN and PPN excitatory input evoked opponent DA responses in TS and DLS. **A)** Experimental strategy to stimulate STN or PPN inputs to SN and monitor DA release in DLS or TS. **B)** Example image shows expression of GRAB_DA_ and location of fiber implant in either DLS or TS; scale, 1 mm. **C)** Optogenetic stimulation of excitatory STN inputs increased DA release in TS (n=6), but in DLS (n=7) led to inhibition of DA followed by a transient increase in DA at stimulus offset inhibited DA responses in DLS compared in inset. **D)** STN-evoked DA responses in TS or **E**) DLS scale with frequency; compared in insets. **F)** Same as in **C)** but with stimulation of PPN inputs, TS (n=9), DLS (n=9). **G)** Same as in **D)** but with stimulation of PPN inputs. **H)** Same as in **E)** but with stimulation of PPN inputs. Sidak’s post hoc, ****p<0.0001.

Stimulation of STN or PPN inputs to SNc evoked strikingly opposite DA responses in TS and DLS (STN: 2-way ANOVA, striatal site: F_1,11_=40.9, p<0.0001, striatal site by time interaction: F_1,11_=40.8, p<0.0001; PPN: striatal site: F_1,16_=36.4, p<0.0001, striatal site by time interaction: F_1,16_ =36.4, p<0.0001) (**Figure 4C, F**). In TS, stimulation of either input elicited an increase in DA release that scaled with frequency (one-way repeated-measures ANOVA, STN 0.25s: F_2,8_=7.5, p=0.02, 1s: F_2,7_=13.6, p=0.004, 4s: F_2,9_=16.9, p=0.001; Friedman’s test, PPN 0.25s: F=20.0, p=0.0002, 1s: F=24.0, p<0.0001, 4s: F=23.2, p<0.0001) (**Figure 4D, G and Figure S3C-D, I-J**) and was observed for all train durations, though the extended 4s train revealed that the evoked DA response peaked and began to decay during the stimulus (**Figure S3A-D and Figure S3G-J**).

In pointed contrast, when we measured DA responses in DLS we found that stimulation of excitatory STN or PPN inputs to SNc evoked a reduction in DA signal below baseline. The DA inhibition was sustained for the duration of the stimulus and was followed by a post-stimulus rebound increase in evoked DA above baseline (**Figure 4C, F**). The inhibition in DA scaled with frequency and was most prominent at higher frequencies (one-way repeated-measures ANOVA or Friedman’s test, STN 0.25s: F_2,9_=7.5, p=0.015, 1s: F_1,11_=12.8 p=0.0005, 4s: F_2,13_=46.8, p<0.0001; PPN 0.25s: F=12.3, p=0.006, 1s: F_1,11_=12.8, p=0.002, 4s: F_2,13_=10.2, p=0.004) (**Figure 4E, H and Figure S3A-B, E-F, G-H, K-L**). Notably, low-frequency (i.e. 5 Hz) optogenetic stimulation of either the STN or PPN increased DLS dopamine levels, an effect we attribute to the summation of individual inhibition-rebound cycles across successive pulses. (**Figure 4E, H and Figure S3F, L**).

Surprised by the inhibition of DA release observed in DLS, we aimed to validate this finding using a complementary approach. We thus used VGLUT2-Cre mice to express ChrimsonR in STN and DAT-Flp to express GCaMP6f in SNc DA neurons; and implanted a fiber in SNc to optogenetically activate STN inputs to SNc, as well as a fiber in TS or DLS to measure calcium fluorescence from DA terminals projecting to these regions (**Figure S3M**). Consistent with the GRAB_DA_ recordings, calcium responses in DA terminals in TS and DLS showed opposite patterns of activity in response to STN stimulation (**Figure S3N-O**). Other hallmarks of the GRAB_DA_ signals were likewise observed in the DA terminal calcium responses including: i) frequency-dependent scaling, ii) transient excitation of TS DA terminals with long train stimulation, iii) post-stimulus rebound excitation in DLS terminals, and iv) summation of inhibition-evoked rebound excitations in the DLS terminals with low frequency stimulation **(Figure S3T-U)**. Combined, the results suggest that excitatory inputs from STN and PPN evoke similar patterns of DA release, but the pattern evoked depends on the DA subcircuit, with increases or decreases in DA release in TS and DLS, respectively. We next aimed to assess whether these responses reflect different activation patterns of Vglut2+ and Anxa1+ DA subtypes.

### Excitatory input from STN and PPN evoke opposite responses in PD-vulnerable and PD-resilient DA subtypes

To record activity of resilient Vglut2+ DA neurons in SNc we used VGLUT2-Cre x DAT-Flp mice with a Cre-plus Flp-dependent AAV (ConFon) to express GCaMP6f in SNc. We also expressed Cre-dependent ChrimsonR in glutamate neurons in either STN or PPN and implanted an optic fiber in SNc. This allowed us to stimulate STN or PPN inputs while simultaneously measuring evoked responses in Vglut2+ DA neurons in SNc (**Figure 5A, E**). To record the activity of Anxa1+ SNc neurons while stimulating STN and PPN inputs we used Anxa1-Cre x VGLUT2-PhiC31. This allowed for Cre-dependent expression (FLEX) of GCaMP8f in SNc to target expression to Anxa1+ neurons, and PhiC31-dependent expression (pSIO) to express ChrimsonR in either STN or PPN. We implanted a fiber in SNc to stimulate inputs, plus a second fiber in DLS to measure calcium activity in Anxa1+ terminals (**Figure 5J, O**).

**Figure 5.**
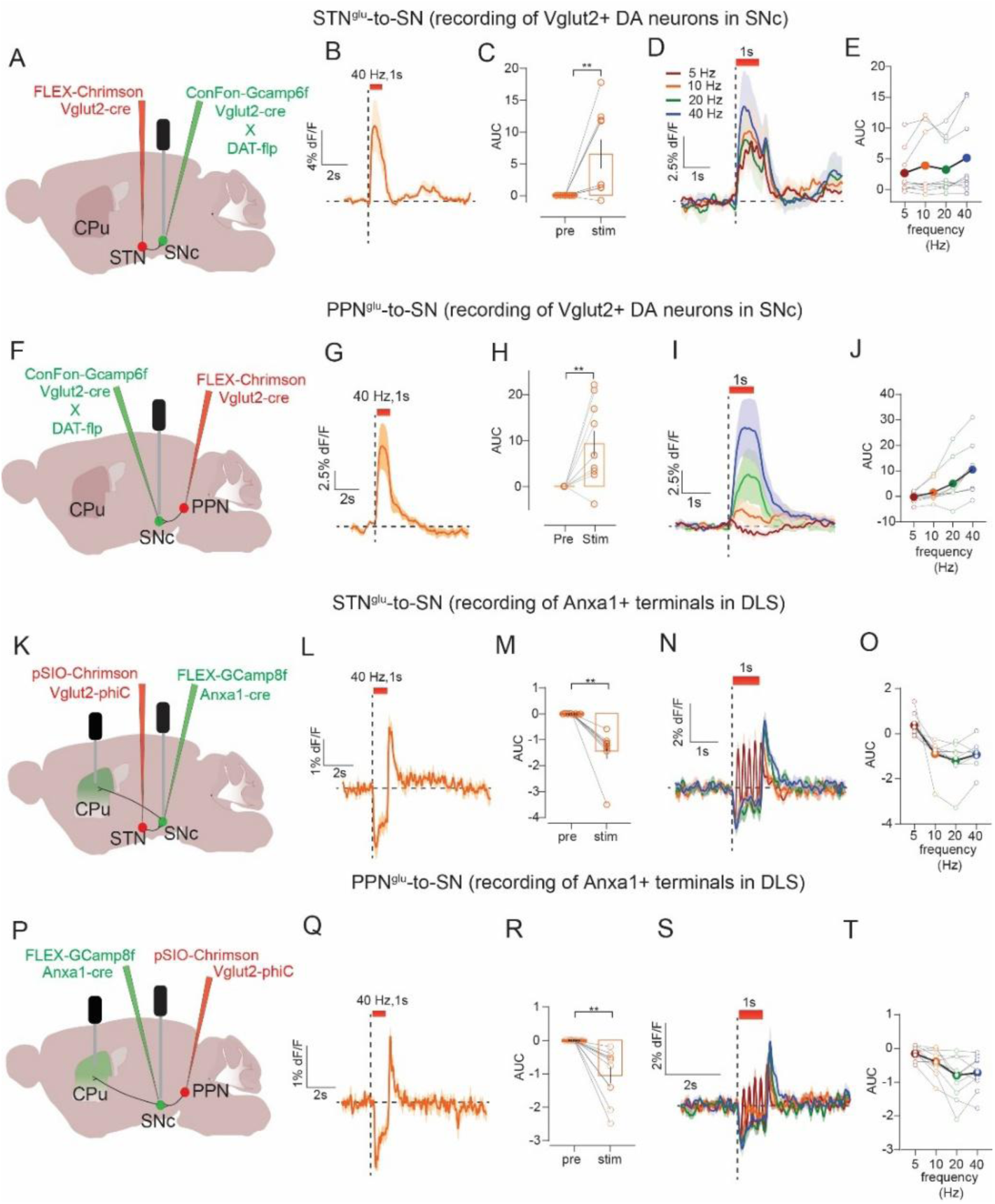
STN and PPN evoke opposite responses in PD-resilient and PD-vulnerable DA subtypes. **A)** Experimental strategy to stimulate STN inputs to SN and record calcium activity in Vglut2-expressing SNc DA neurons. **B)** Optogenetic stimulation of STN inputs (n=10) increased activity in Vglut2+ DA neurons, **C**) during 1s stimulation compared to pre-stimulus period. **D**) Comparing across frequency **E**) did not reveal a significant effect of frequency. **F**) Same as **A)** but stimulation of PPN inputs, which **G**) also increased activity in Vglut2+ DA neurons (n=10) **H**) compared to pre-stimulus period. **I)** Comparing across frequency **J**) revealed a frequency-dependent increase. **K)** Strategy to stimulate STN inputs to SN and record calcium activity in Anxa1+ neurons in SNc or from their DLS terminals. **L)** Optogenetic stimulation of STN inputs (n=8) inhibited activity in Anxa1+ terminals in DLS but led to a transient increase in their activity at stimulus offset, **M**) inhibition compared to pre-stimulus period. **N**) Comparing across frequency **O**) did not reveal a significant effect of frequency. **P**) Same as K) but stimulation of PPN inputs, which **Q**) also led to inhibition of activity in Anxa1+ terminals in DLS with a transient post-stimulus increase in activity (n=10), **R**) compared to pre-stimulus period. **S)** Comparing across frequency **T**) revealed a frequency-dependent inhibition in activity. Paired t-test or Wilcoxon, **p<0.01.

Stimulation of either STN or PPN inputs evoked an increase in the activity of Vglut2+ DA neurons in SNc (Wilcoxon’s test or paired t-test, STN: 0.25s: W=41.0, p=0.012, 1s: W=43.0, p=0.008, 4s: W=-5.0, p=0.82; PPN: 0.25s: t=2.9, p=0.02, 1s: t=3.4, p=0.008, 4s: t=3.0, p=0.01) (**Figure 5C, H and Figure S4A, 4D**). Consistent with results from TS GRAB_DA_ recordings, increases in activity scaled with frequency for short-duration STN stimulations and all durations tested for PPN (Friedman’s test STN: 0.25s: F=15.5, p=0.001, 1s: F=5.9, p=0.11, 4s: F=7.0, p=0.07; PPN: 0.25s: F=18.8, p=0.0003, 1s: F=24.8, p<0.0001, 4s: F=20.2, p=0.0001), and long stimulus trains (i.e., 4s) evoked a transient increase followed by a decay. In a few mice for both STN and PPN inputs, we noticed that the decay dropped below baseline, resulting in a net inhibition of activity as assessed by an area under the curve (AUC) measurement that aggregates the change in signal during the stimulus train (**Figure S4A, D**).

In DLS terminals from SNc Anxa1+ neurons, stimulation of both STN and PPN inputs produced inhibition followed by apparent rebound at stimulus offset (paired t-test or Wilcoxon’s test, STN: 0.25s: t=6.4, p=0.0004, 1s: W=-36.0, p=0.005, 4s: Wilcoxon’s test, W=-36.0, p=0.008; PPN: 0.25s: t=3.9, p=0.004, 1s: W=-55.0, p=0.002, 4s: t=3.8, p=0.004) (**Figure 5L-M, Q-R and Figure S4B, E**). The inhibition again scaled with frequency for shorter-duration STN stimulation and all durations tested for PPN (Friedman’s test or one-way repeated-measures ANOVA, STN: 0.25s: F=18.0, p=0.0003, 1s: F=15.5, p=0.015, 4s: F=15.3, p=0.001; PPN: 0.25s: F=11.5, p=0.0002, 1s: F_2,15_=11.0, p=0.006, 4s: F_2,18_=7.2, p=0.002), and low-frequency stimulation (i.e., 5 Hz) produced a small dip and rebound with each laser pulse. Finally, we also made recordings from Anxa1+ SNc cell bodies in the same animals. We prioritized analysis of Anxa1+ SNc terminals in DLS because cell body signals were only detected in a subset of mice (due to low GCaMP expression or off-target fiber implants). However, analysis of that subset revealed the same pattern of responses including stimulation-induced inhibition of activity (paired t-test, STN: 0.25s: t=3.1, p=0.035, 1s: t=4.0, p=0.016, 4s: t=7.2, p=0.002; PPN: 0.25s: t=4.8, p=0.018, 1s: t=8.4, p=0.004, 4s: t=3.3, p=0.044) followed by post-stimulus rebound activation (**Figure S4C, F**). In sum, multiple in vivo recording approaches point to the same conclusions, that excitatory inputs from STN and PPN led to activation of PD-resilient Vglut2+ DA neurons but inhibition of PD-vulnerable Anxa1+ DA neurons.

## Discussion

*Anxa1* was recently identified in a subset of mouse SNc DA neurons and several lines of evidence indicate it could serve as a new marker of PD-vulnerable DA neurons. In mice, *Anxa1* is expressed within a subset of the *Sox6*+ and *Aldh1a1*+ DA neurons (Azcorra et al. 2023; Gaertner et al. 2025) markers already known to label vulnerable DA neurons in human PD (Liu et al. 2014; Pereira Luppi et al. 2021). *Anxa1*+ DA neurons localize to ventral SNc and project to DLS, where their activity correlates with locomotor acceleration (Azcorra et al. 2023). Finally, *Anxa1*+ DA neurons were recently shown to be preferentially vulnerable in mouse models of PD (Fushiki et al. 2024; Mantas et al. 2024). However, there has been little evidence that *ANXA1* is expressed in human DA neurons or whether its expression is altered in PD (Knott et al. 2000; Kamath et al. 2022). Thus, before using *Anxa1* as a marker of PD-vulnerable neurons in our mouse studies we investigated its expression in human PD.

We found that *ANXA1* mRNA was present in a subset of human SNc DA neurons and was de-enriched among surviving DA neurons in PD, indicating it is a marker of PD-vulnerable neurons. Approximately 8% of SNc DA neurons expressed *ANXA1* in age-matched non-PD control cases, but *ANXA1*+ DA neurons were virtually non-existent in the majority of PD cases. Examination of *ANXA1* expression patterns suggest localization to DA neurons within SNc subregions similar to *ALDH1A1*+ and ventral to *CALB1*+ DA neurons. Comparing these three DA subtype markers within the same cases suggest that *ANXA1* may be most strongly de-enriched in PD, and thus most sensitive to PD-related neurodegeneration, and that loss of these DA neurons may thus contribute to the earliest symptoms associated with nigrostriatal DA dysfunction in PD. However, future human and animal studies will be needed to further support or reject this hypothesis.

Having established *ANXA1* as a marker of PD-vulnerable DA neurons we sought to compare how afferent inputs to SN could generate different activity patterns within different DA populations. This question is important because distinct DA subtypes and subcircuits have divergent patterns of activity that appear to contribute to diverse roles for DA in the regulation of movement and other DA-dependent behaviors. We focused on *in vivo* measurements of two DA subpopulations that are largely non-overlapping: the *Anxa1*+ PD-vulnerable DLS-projecting population of DA neurons that increase activity with locomotor acceleration (Azcorra et al. 2023), compared to the Vglut2+ population of DA neurons that we and others have shown are preserved in PD animal models and resilient to neurodegeneration in human PD (Mendez et al. 2008; Shen et al. 2018; Steinkellner et al. 2018; Steinkellner et al. 2022; Buck et al. 2021; Fushiki et al. 2024). Vglut2+ PD-resilient DA neurons mainly localize to lateral SNc, project densely to TS, and their activity increases with locomotor deceleration (Azcorra et al. 2023). Thus Vglut2+ and *Anxa1*+ DA populations have divergent molecular, physiological, anatomical, and pathological features that make them an attractive choice for comparative assessment.

To understand how distinct DA activity patterns may be generated by excitatory afferents we selected two targets for optogenetic stimulation: STN and PPN. When we first measured STN-or PPN-evoked responses from a non-selective population of DAT-expressing DA neurons, observed responses were highly variable between subjects and displayed complex temporal dynamics that included apparent excitations and inhibitions. Subsequent experiments are consistent with the hypothesis that these complex responses represented summations of distinct DA subtype responses, demonstrating the crucial importance of considering DA neuron heterogeneity.

DA recordings from TS or calcium recordings from Vglut2+ DA neurons in SNc revealed a more straightforward response; excitatory inputs from either STN or PPN evoked an increase in DA activity. This increase is consistent with prior work suggesting that SNc DA neurons receive direct monosynaptic input from glutamate neurons in STN and PPN (Lavoie & Parent 1994; Charara et al. 1996; Watabe-Uchida et al. 2012; Menegas et al. 2015), and that activations of STN or PPN can evoke DA responses, especially in TS (Scarnati et al. 1984; Smith & Grace 1992; Iribe et al. 1999; Lokwan et al. 1999; Todd et al. 2022). The increase in DA activity largely scaled with increasing frequency and train duration, though for trains longer than 1s the response typically peaked and began to decay during the stimulus train.

In striking contrast, DA recordings from DLS or calcium recordings from *Anxa1*+ SNc neurons showed pronounced inhibitory responses. DA release and calcium activity were both inhibited when excitatory inputs from STN or PPN were stimulated. This inhibition was sustained for the duration of the stimulus train, but was followed by a pronounced increase in DA activity above baseline upon stimulus offset, consistent with a post-stimulus rebound activation of DA neurons. On the one hand this result is surprising because STN and PPN provide broad monosynaptic excitatory input to SNc DA neurons (Watabe-Uchida et al. 2012; Galtieri et al. 2017), which likely includes DLS-projecting *Anxa1*+ DA neurons in ventral SNc. However, these ventral SNc DA neurons also receive GABAergic input from SNr and other nuclei downstream of STN and PPN activity (Fushiki et al. 2024; Wu *et al*., 2019). Indeed, SNr GABA neurons are a canonical target of STN and PPN and we observed their large, sustained, and frequency-dependent activation in response to STN or PPN stimulation. Moreover, ventral SNc DA neurons, including *Anxa1*+ DA neurons, make unique dendritic structures that extend ventrally into SNr, so-called striosome-dendron bouquets, composed of intertwined DA dendrites plus GABAergic and other afferents (Crittenden et al. 2016; Mantas et al. 2024). We speculate that these specializations serve as anatomical substrates that constrain the ability of excitatory inputs to activate PD-vulnerable *Anxa1*+ DA neurons. One possibility is that, *in vivo*, excitatory inputs from STN or PPN only drive *Anxa1*+ DA neurons when there is coincident inhibition of SNr GABA neurons, for example from pallidum or striatum. Indeed, different populations of striatonigral neurons were recently shown to have distinct effects on locomotion and dopamine release, potentially through differential inhibitory and disinhibitory effects on SNc DA subtypes (Dong et al. 2025). Another possibility is that heterogeneous subpopulations of STN or PPN neurons (Wallén-Mackenzie et al. 2020; Masini & Kiehn 2022; Goñi-Erro et al. 2023) selectively innervate distinct populations of SNc DA and SNr GABA neurons. Future experiments may dissect these and other non-mutually exclusive possibilities.

A second feature of *Anxa1*+ DA neuron responses to excitatory input is the pronounced post-stimulus rebound activation. This *in vivo* response is reminiscent of prior *ex vivo* slice studies that reported rebound activation following direct hyperpolarization via current injection or by activation of GABA_B_ receptors on DA neurons. These studies demonstrated that it was only a subset of DA neurons, ventrally located or *Aldh1a1*+, that displayed this rebound signature (Fiorillo et al. 2013; Evans et al. 2020; Beaver & Evans 2025). The inhibition-evoked rebound excitation appeared to represent an intrinsic property of these DA subtypes, dependent on strong expression of hyperpolarization-activated cyclic nucleotide-gated channels and T-type calcium channels (Evans et al. 2017; Evans 2022). Moreover, other recordings from DA neurons *in vivo* have shown this inhibition-excitation, or pause-rebound, motif. For example, subsets of DA neurons show a pause-rebound response to aversive or other salient stimuli in both rodents and primates (Fiorillo & Williams 1998; Brischoux et al. 2009; Wang & Tsien 2011; Fiorillo et al. 2013) or following stimulation of inputs to SN from PPN or striatal GABA neurons or glutamate inputs from superior colliculus (Gut et al. 2022; Poisson et al. 2024; Dong et al. 2025). An exciting direction for future work will be to test whether this inhibition-evoked excitation represents a physiologically and behaviorally relevant mechanism for evoking DA release in DLS.

Our findings may also have important implications for understanding how deep brain stimulation (DBS) differentially affects vulnerable and resilient DA circuits. STN DBS is an important therapeutic approach used to treat PD symptoms. It remains debatable whether high-frequency DBS has excitatory or inhibitory effects on STN output (Herrington et al. 2016; Lozano et al. 2019; Alosaimi et al. 2022; Schor et al. 2022), but our results suggest that the impact of changes in STN output are likely to have diverse effects on DA subtypes. For example, STN DBS could produce prokinetic effects on behavior by reducing activity in resilient Vglut2+ DA neurons that have been associated with deceleration. Alternatively STN DBS could produce prokinetic effects by disinhibiting surviving *Anxa1*+ DA neurons that have been associated with acceleration. Testing these ideas in animal models or human PD could allow for optimization of DBS procedures that maximize the utility of residual DA in discrete subcircuits.

One limitation of our approach is the reliance on stimulus-evoked responses driven by optogenetic activation of global populations of excitatory neurons in STN or PPN. This approach leads to broad synchronized activation of excitatory inputs to SN and could result in supraphysiological responses and erroneous inferences. However, our use of input-output curves to compare across stimulation patterns ranging from as little as a single pulse provides some assurance that the effects observed are not a function of a particular non-physiological pattern of STN or PPN activity. Moreover, we note that this challenge is inherent to most *ex vivo* or *in vivo* studies that aim to assess functional connectivity and make inferences therefrom. We also note that the maximum frequency we tested (40 Hz) is well below frequencies commonly used for STN DBS, typically greater than 100 Hz. Our results nevertheless reveal that a given stimulation parameter can drive opposing effects on different DA subtypes. Another potential limitation is our use of calcium and DA sensors for photometric recordings of bulk changes in cell-type activity. Though these approaches provide several advantages, they obscure heterogeneity within a specified cell type and offer temporal resolution on the order of tens to hundreds of ms, 1 or 2 orders of magnitude slower than single-unit electrophysiological recordings. Thus stimulus-evoked dynamics that occur more quickly may not be well represented in our dataset. Additional work that aims to assess the effects of manipulating STN or PPN subtypes or applying technologies for higher resolution temporo-spatial patterning of stimulation may yield results that support or alter present conclusions that are based on currently available data.

In summary, our findings reveal that excitatory inputs from the STN and PPN differentially modulate PD-resilient and PD-vulnerable DA subtypes in vivo. Specifically, we demonstrate that Vglut2+ DA neurons, resilient in both mouse models and human PD, are activated by these inputs; while Anxa1+ DA neurons, which we show are highly vulnerable in PD, are inhibited followed by rebound excitation. These results suggest that DA subtypes are embedded in distinct functional circuits that shape their activity patterns, behavioral functions, and perhaps their susceptibility to neurodegeneration. Moreover, the subtype-specific responses to STN and PPN input raise the possibility that neuromodulatory interventions, including DBS, may differentially impact residual DA subcircuits depending on their anatomical and molecular identity. Identifying and understanding the organizational logic of these subtype-specific circuits could thus lead to more targeted and effective strategies for restoring motor function in PD.

## Author Contributions

Conceptualization, L.C.H., N.G.H., and T.S.H.; Methodology L.C.H., N.G.H., E.B.L., B.L., F.C., R.A and T.S.H. Investigation L.C.H., N.G.H., G.J.K., L.L., M.X. and M.S.; Formal analysis L.C.H., N.G.H. and M.S.; Writing – original draft, L.C.H., N.G.H., and T.S.H.; review and editing, L.C.H., N.G.H., G.J.K., L.L., M.X., M.S E.B.L., B.L., F.C., R.A and T.S.H.; funding acquisition T.S.H.

## Acknowledgments

This work was funded by the Aligning Science Across Parkinson’s (ASAP) through the Michael J. Fox foundation for Parkinson’s Research (MJFF) and the National Institutes of Health (R01 NS138598). MJFF administers the grant (ASAP-020600) on behalf of ASAP and itself. The collection of human postmortem tissue provided by the Center for Neurodegenerative Disease research at University of Pennsylvania (CNDR) is supported by the funding from the National Institute on Aging (P01AG084497). For the purpose of open access, the author has applied a CC-BY 4.0 public copyright license to the Author Accepted Manuscript (AAM) version arising from this submission.

## Materials and Methods

### Chromogenic *in-situ* hybridization assay

Human tissue blocks obtained from the National Institutes of Helath NeuroBioBank (through the University of Miami, Miller School of Medicine, Brain Endowment Bank) were sectioned at 5-um using a microtome (Microm) and mounted on glass slides (Super frost Plus, Fisher Scientific, 1255015). Samples obtained from the University of Pennsylvania were processed similarly and shipped to the University of California, San Diego (UCSD). Slide-mounted sections were stored at room temperature with dessicant. The slides were baked at 60°C for 1h followed by deparafinization with xylenes twice for 5 min each. Sections were rinsed twice with 100% ethanol (EtOH) and air-dried 5 min. Subsequently, sections were pre-treated with hydrogen peroxide (ACD, 322335) for 10 min followed by an incubation in target retrieval solution (ACD, 322001) for 15 min at 97°C. Sections were rinsed with double distilled water (ddH_2_O) and dried at 60°C on a heat block. A hydrophobic barrier was drawn around the samples with a pen (Scientific Device laboratory, 9804-02) followed by an incubation with ProteasePlus (ACD, 322331) at 40°C for 30 min. Sections were hybridized for 2h at 40°C using probes targeting *TH* (ACD, 441651-C2) and either *CALB1* (ACD, 422161), *ALDH1A1* (ACD, 310151), or *ANXA1* (ACD,465411). Sections were stored in saline-sodium citrate (SSC 5X, Thermo scientific, J60839.K2) buffer overnight. The next day, sections underwent 10 amplification cycles using (ACD, 322500), counterstained with Hematoxylin (Sigma, GHS132-1L) for 30s and dipped in tap water. The sections were dried at 60°C for 20 min and coverslipped with Vectashield mounting medium (H-500). Images were acquired using brightfield imaging (Axioscan Z1, Zeiss) at 20X magnification. The SNc borders were defined according to the brain atlas published by Couloumb et al (Coulombe et al. 2021). Quantification was done manually using Zen Blue software (version 3.1, Zeiss) and the distribution maps of DA subtypes were generated with Adobe Illustrator (Version 27).

### Immunofluorescence on human tissue

Formalin-fixed paraffin embedded 5-µm-thick sections of the SNpc were used to perform the immunofluorescence. Sections were obtained through the Parkinson’s UK Brain Bank under the REC of the CHU de Québec ethical approval 2016-2564. Sections were placed on a slide warmer at 65°C for 20 min to remove the paraffin, followed by defatting in Citrisolv (Fisher Scientific, cat# 22143975) and rehydratation in descending ethanol solution baths. Sections were rinsed with ddH_2_O and washed in phosphate-buffered saline 1X (PBS, Fisher Scientific, cat# BP399) with 0.1% Tween (Tween-20, Fisher Scientific, cat# BP337) (PBS-T) before an antigen retrieval was performed with Tris-EDTA buffer (Quattro sol II, Diagnostic Biosystems, cat# K086-RUO) pH 9.0 at 95°C for 40 min. After 20 min cooling at room temperature (RT) in the buffer, sections were washed with PBS-T and then immersed in blocking solution (1% bovine serum albumin [BSA, BioShop Canada, cat# ALB001], 1% Triton X-100 [MilliporeSigma, cat# T8787], 10% Normal donkey serum [NDS, MilliporeSigma, cat# D9663] in PBS 1X) for one hour at RT. After additional washes, sections were incubated overnight at 4°C with a mix of primary antibodies in blocking solution; goat anti-ALDH1a1 (1/250; R&D BioTechne, cat# AF5869) and mouse anti-TH, 1/500; MilliporeSigma, cat# MAB318). The following day, sections were washed with PBST and incubated with secondary antibodies diluted in blocking solution; donkey anti-goat Alexa Fluor 647 (1/500; Invitrogen, cat# A21447) and donkey anti-mouse Alexa Fluor 488 (1/500; Invitrogen, cat# A21202) for 2.5 hours at RT. After subsequent washes in PBS 1X, sections were incubated with DAPI (1/5000, DAPI 5mg/mL, Invitrogen, cat# D3571) for 10 min, washed in PBS 1X and autofluorescence was quenched using a solution of 0.1% Sudan Black (MilliporeSigma, cat# 199664) in 70% ethanol. Sections were washed in PBS to remove excess Sudan Black, rinsed in ddH_2_O and coversliped with Fluoromount-G (Invitrogen, cat# 00-4958-02). After drying scans of the entire section were acquired with an AxioScan.Z1 (Zeiss) using Zen software (ZEN 3.1, Zeiss). Cell counting was performed with Stereo Investigator software (Version 2019, MicroBrightField Bioscience) and mappings were retrieved with NeuroExplorer software (Version 2016, MicroBrightField Bioscience).

### Animals

All mouse experiments were performed in accordance with protocols approved by UCSD Institutional Animal Care and Use Committee. Male and female mice were bred at UCSD and group-housed on a 12-hour light/dark cycle, with food and water *ad libitum*. VGLUT2-Cre (*Slc17a6^tm2(cre)Lowl^*, RRID:IMSR_JAX:016963) and VGAT-Flp (*Slc32a1^tm1.1(flpo)Hze^*, RRID:IMSR_JAX:029591) mice were obtained from the Jackson Laboratory and maintained backcrossed on to C57BL/6J (RRID: IMSR_JAX:000664). The DAT-Flp (*Slc6a3^em1(flpo)Hbat^*, RRID:IMSR_JAX:035436) were generously provided by Dr. Helen Bateup (UC Berkeley). VGLUT2-PhiC31 (Jeong et al. 2024) mice were generously provided by Dr. Lim (UC San Diego). DAT-Flp (*Slc6a3^em1(flpo)NW^* Jax and RRID) and Anxa1-Cre (RRID:MMRRC_069931-MU) were generously provided by Dr.Awatramani (Northwestern).

### Stereotactic surgeries

Mice >6 weeks old were anesthetized with isofluorane (4% for induction and 1-2% for maintenance), and provided with pre- and post-surgical analgesic carprofen (5 mg/kg i.p). For calcium recordings in SNr or SNc VGLUT2-Cre x VGAT-Flp or VGLUT2-Cre x DAT-Flp mice were injected with 300 nl of AAV1-Ef1a-fDIO-GCaMP6f (4e12 vg/mL, Addgene 1283125) in SNr (AP −3.3, ML +/−1.3, DV −4.6; all coordinates relative to Bregma) or SNc (AP −3.2, ML +/−1.3, DV −4.3). For optogenetic stimulation, 150-200 nl of AAV5-Syn-FLEX-ChrimsonR-tdTomato (4 – 8.5e12 vg/ml, Addgene 62723) was injected in either STN (AP −2.00, ML +/−1.6, DV −4.5) or PPN (AP −4.48, ML +/− 1.1, DV −3.75). A fiber (400-um core, 0.39 NA, 6-mm length, 1.25-mm diameter black ceramic ferrule, RWD) was implanted in either SNr (AP −3.3, ML +/−1.4, DV −4.4) or SNc (AP −3.2, ML +/−1.4, DV −4.1).

For monitoring DA release in striatal subregions VGLUT2-Cre mice were injected with AAV5-Syn-FLEX-ChrimsonR-tdTomato in either STN or PPN as above. And 300 nl of AAV9-EF1a-GRAB_DA2m_ (2e12 vg/mL, Addgene: 140553) was injected in either DLS (AP +0.5, ML +/−2.25, DV −2.95) or TS (AP −1.25, ML +/−2.9, DV −3.45). A fiber was implanted in SNc (200-um core, 0.39 NA, L = 6 mm, 1.25-mm diameter white ceramic ferrule, RWD) and a second fiber (400-um core, 0.39 NA, L = 4 mm, 1.25-mm black ceramic ferrule, RWD) was implanted in DLS (AP +0.5, ML +/− 2.25, DV −2.75) or TS (AP −1.25, ML +/− 2.9, DV −3.25).

For recording from Vglut2+ DA neurons in SNc, VGLUT2-Cre x DAT-Flp animals were injected with 300 nl of AAV8-Ef1a-ConFon-GCaMP6f (2.3e12 vg/ml, Addgene: 137122) in SNc (AP −3.20, ML +/−1.5, DV −4.20), and AAV5-Syn-FLEX-ChrimsonR-TdTomato was injected in STN or PPN as above. A fiber (400-um core, 0.39 NA, L = 6 mm, 1.25 mm black ceramic ferrule, RWD) was implanted in SNc (AP −3.20, ML +/−1.60, DV −4.00). For recording from Anxa1+ neurons and terminals projecting to DLS, VGLUT2-PhiC x Anxa1-Cre mice were injected with 150-200 nl of AAVDJ-Ef1a-pSIO-ChrimsonR-tdTomato (1e13 vg/ml, Vectorbuilder) in STN or PPN at the same coordinates as above. And injected with 300 nl of AAV5-Syn-FLEX-GCaMP8f (1e13 vg/ml, Addgene: 162379) in SNc (AP −3.20, ML +/−1.30, DV −4.30). Optic fibers were implanted in both SNc and DLS (400-um core, 0.39 NA, L = 6 or 4 mm, 1.25-mm black ceramic ferrule, RWD).

### Fiber photometry recordings with optogenetic stimulation

Signal processing and acquisition hardware (RZ5P or RZ10x; Tucker-Davis Technologies, USA) was used to control 465 nm and 405 nm LEDs for excitation of the sensor’s fluorescence signal or isosbestic channel, respectively modulated at 210 Hz and 330 Hz, filtered and combined by a fluorescence mini cube (Doric Lenses, USA), and delivered through a 400-µm core, 0.57 NA, low-autofluorescence fiber optic patch cable coupled to a pigtailed rotary joint (Doric). Event timestamps were digitized in Synapse Software (Tucker Davis Technologies) via TTL signals issued from AnyMaze (Stoelting) to synchronize with optogenetic laser onset. LED excitation power used for fiber photometry recording (<0.2 mW) was an order of magnitude less than that used for optogenetic laser stimulation (5mW).

Optogenetic stimulations were delivered by a red laser (Shanghai laser, 635nm) at 5 mW power. The mice received 40 Hz stimulations at various durations (i.e. 250 ms, 1 s and 4 s), 5 trials for each duration, with an inter-stimulation interval (ISI) of 10 s, and trials within session and subject were averaged. In subsequent recording sessions, mice received optogenetic stimulation at varying frequencies (5 Hz, 10 Hz, 20 Hz and 40 Hz), 5 trials for each, with an ISI of 20 s. For a given session block, the stimulation durations were again either 250 ms, 1 s, or 4 s, with the order of stimulation duration blocks (ascending or descending) counterbalanced across animals.

### Fiber photometry signal extraction and analysis

Analysis of recorded signals was performed using custom MATLAB code (Mathworks, RRID: with the signal processing, statstics, and curve fitting toolboxes). Fluorescent traces were demodulated offline, filtered with a 6 Hz lowpass zero-phase digital filter, and downsampled to 100 Hz. The isosbestic channel from the 405-nm excitation LED was fit with a biexponential curve, which was fit using robust linear regression to the signal channel from the 465-nm excitation LED. This fitted curve was then used to calculate deltaF over F (dF/F). Traces were aligned to time of each stimulation onset, and the mean dF/F during a 1-s pre-stimulation baseline period was subtracted. The area under the curve (AUC) was calculated within a given time window using trapezoidal integration for the specified stimulation durations.

### Histology

Mice were deeply anesthetized with sodium pentobarbital (200 mg/kg; i.p.; VetOne) and transcardially perfused with 30-40 mL of phosphate buffer saline (PBS) followed by 60-70 mL of 4% paraformaldehyde (PFA) at ∼ 7mL/min. Brains were extracted and incubated in PFA overnight at 4°C followed by cryoprotection in 30% sucrose for 48h. Brains were flash frozen in isopentane and stored at −80°C. Brains were sectioned at 50 microns in a cryostat (Leica) and stored in PBS with 0.01% sodium azide at 4°C. For immunostaining, sections were washed 3 times for 5 min with PBS. Sections were rinsed 3 times for 5 min with PBS containing 0.2% triton X-100 (PBST) followed by blocking for 1h with PBST containing 4% normal donkey serum (Jackson Immunoresearch) at RT. Sections from SN and striatum were subsequently incubated in sheep anti-TH (1:2000, Pelfreeze P60101-0), rabbit anti-DsRed (1:2000, clonetch 632496), and chicken anti-GFP (1:2000, invitrogen A10262) at 4°C overnight. Some sections containing PPN were incubated with goat anti-ChAT (1:400, invitrogen AB144P) at room temperature overnight. The next day, sections were washed 3 times for 5 min with PBS followed by incubation with secondary antibodies conjugated with Alexa-647 (Donkey anti-sheep, 1:400), Alexa-594 (Donkey anti-rabbit, 1:400), Alexa-488 (Donkey anti-chiken, 1:400), and/or Alexa 647 (Donkey anti-goat, 1:400) (Jackson Immunoresearch) for 2h at RT. The sections were rinsed 3 times with PBS 5 min each and subsequently incubated in PBS containing DAPI (1:1000, Sigma) for 5 min. Sections were washed 3 times with PBS 1X 5 min each, mounted onto glass slides, and coverslipped with Fluoromount-G mounting medium (Southern Biotech).

Images were acquired using a widefield epifluorescence microscope (Zeiss Axioscan Z1). Tiled images were taken at 10X with appropriate filters and using identical settings across slides. Approximately 3-4 sections through the rostral-caudal extent of DLS or TS, 3-4 sections of SNc, 4-8 sections of STN, and/or 3-4 sections of PPN were imaged. ChrimsonR-tdTomato, GCaMP, and GRAB_DA_ expression and optic fiber placement were mapped onto corresponding coronal sections in the Paxinos Mouse Brain Atlas using Adobe Illustrator (version 27). We excluded 26 animals based on 3 exclusion criteria: (1) We observed ChrimsonR-TdTomato in SNc cell bodies in 2 animals, (2) the fiber placement was too dorsal to the intended target (> 500 um) or otherwise misplaced in 9 animals and (3) There was an absence of viral expression in 15 animals.

### Statistical analysis

Data was analyzed with GraphPad Prism (v10). We assessed the distribution with Shapiro-Wilk test. If normally distributed, the data were analyzed by either using Student’s t-test, one-way ANOVA, or 2-way ANOVA followed by Sidak’s post-hoc multiple comparisons. Else the data were analyzed by either using Mann-Whitney, Wilcoxon test, or Friedman test. All data are represented as mean ± standard error of the mean (SEM) and/or as individual points. Statistical details experiments, including statistical tests and sample size can be found in results and figure legends.

---

**Supplemental figure 1 (related to figure 1).**
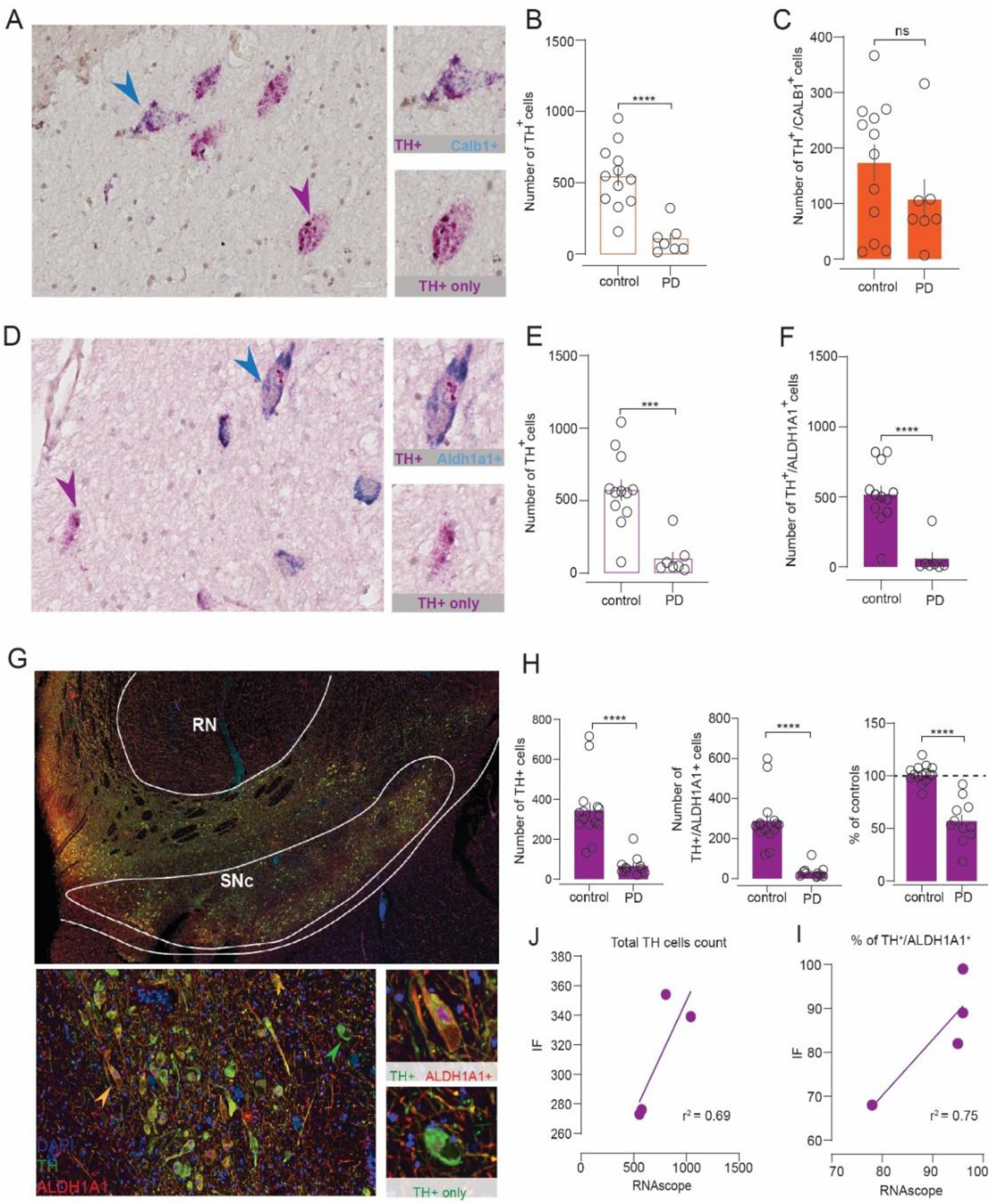
Quantification of *CALB1* and *ALDH1A1* populations in SNc from same cases as described for *ANXA1*. **A)** Example images showing SNc DA neurons expressing *TH* alone (purple arrow) or both *TH* and *CALB1* (blue arrow). Scale: 50 μm (top), 10 μm (bottom). **B)** *TH*+ neuron counts were significantly reduced in PD cases (n=7) compared to controls. **C)** *TH*+/*CALB1*+ neuron counts were unchanged between groups. **D)** Same as A), showing *TH* and *ALDH1A1* labeling. **E)** Same analysis as **B)** for *ALDH1A1*. **F)** *TH*^+^/*ALDH1A1*^+^ counts are significantly reduced in PD. **G)** Representative immunofluorescence (IF) images at low and high magnification showing *DAPI* (blue), *TH* (green), and *ALDH1A1* (red) in SNc from a control case. **H**) Quantification of IF showing significant decrease in TH+, *TH^+^/ALDH1A1^+^*, and % *TH+/ALDH1A1+* showing de-enrichment in PD (n=11) relative to control (n=15). **I)** A subset of cases (n=4) were processed by both IF and RNAscope, showing correlated TH+ counts and **J)** fraction of *TH+* cells expressing *ALDH1A1*. Unpaired t-test or Mann-Whitney, ***p<0.001, ****p<0.0001.

**Supplemental Figure 2 (related to Figures 2-5, S3-S4).**
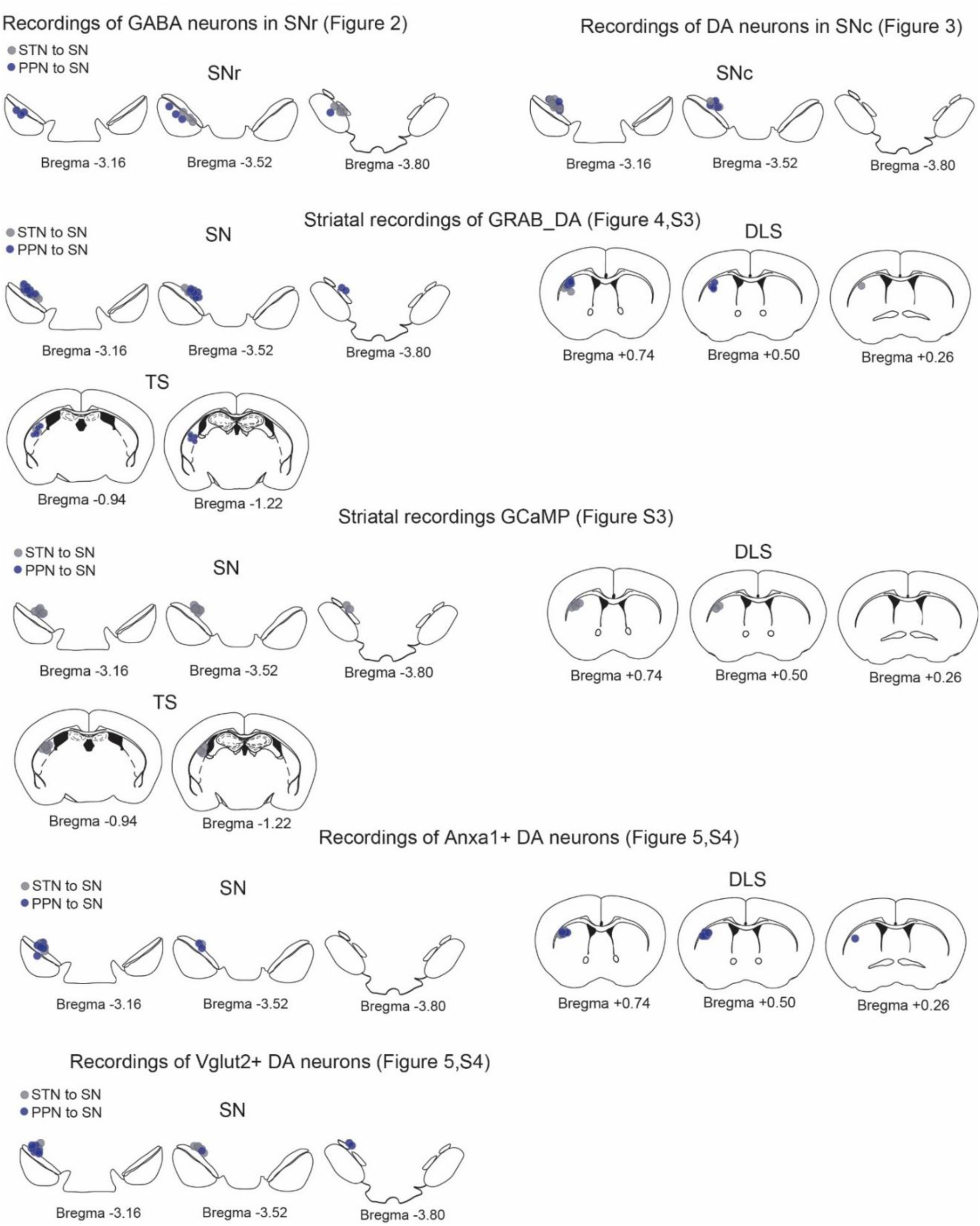
Maps showing fiber placement for each animal included in Figures 2-5 and S3-S4 in SN, DLS, or TS. All placements depicted on the left hemisphere here for ease of comparison. Actual implants were counterbalanced between the left vs. right hemispheres across mice.

**Supplemental figure 3 (related to figure 4).**
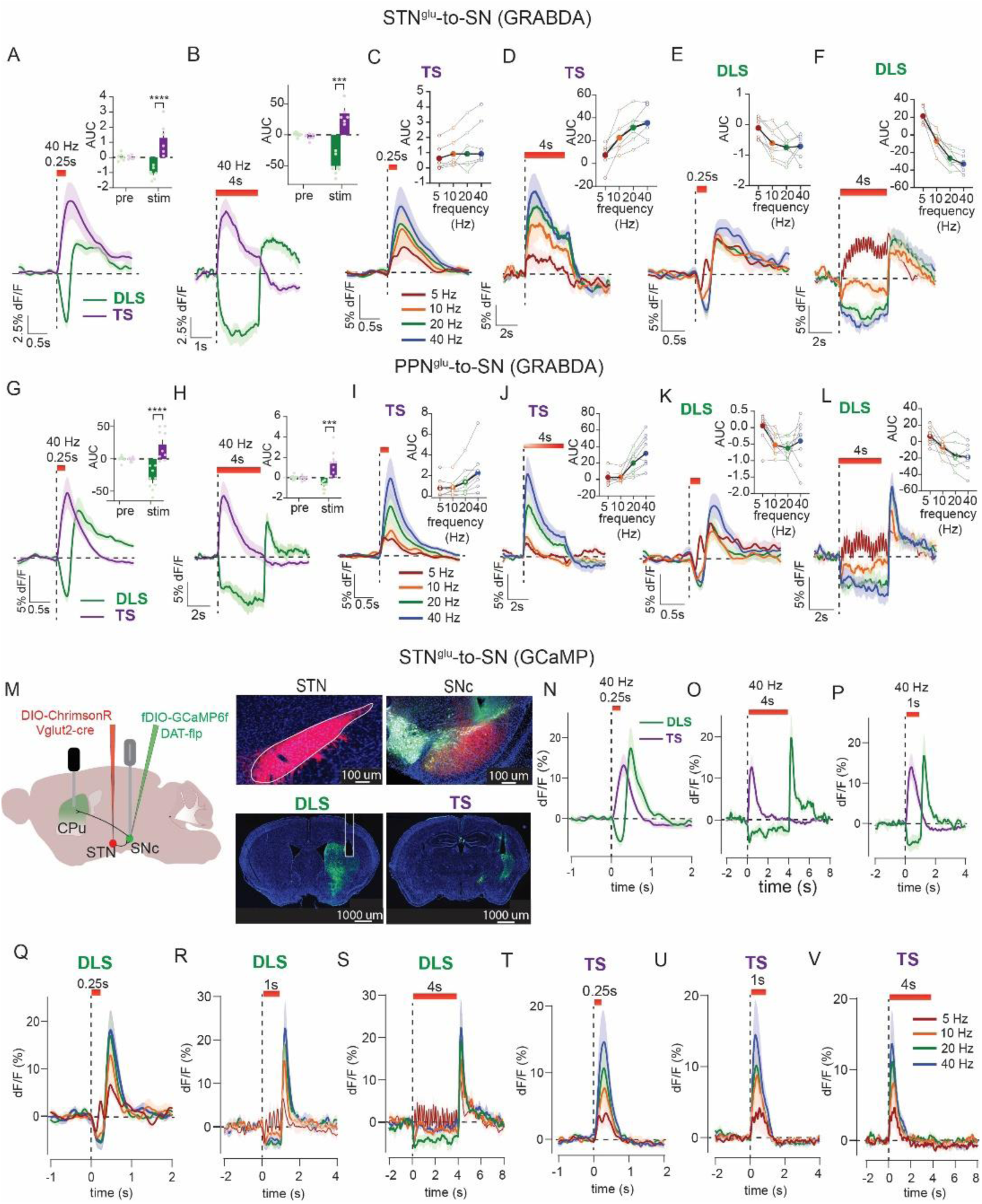
Comparing DA responses to STN and PPN stimulation across additional parameters. **A)** Optogenetic stimulation of STN inputs for 0.25 s or **B**) 4 s evoked DA release measured by GRAB_DA_ in TS (n=6) but led to suppression of DA in DLS (n=7). **C-F)** Varying frequency of STN stimulation with 0.25 s or 4 s trains produced frequency-dependent increases in TS or decreases in DLS. **G-L)** Same as **A-F)** but with stimulation of PPN inputs (TS and DLS: n=9). **I–L)** PPN stimulation produced a frequency-dependent increase in TS, and inhibition in DLS. **M)** Experimental strategy to stimulate STN inputs to SN and record the activity of SNc DA terminals in TS and DLS with GCaMP6f; histology showing ChrimsonR:TdTomato expression in STN neurons and SN terminals (red), plus GCaMP expression in SNc DA neurons and their terminals in DLS or TS, sections counter-stained with DAPI (blue) and in SNc with TH (white); scale 1 mm. **N-V)** Stimulation of STN inputs to SN and recordings of activity in DA terminals in TS (n=9) or DLS (n=9). Sidak’s post hoc, ***p<0.001, ****p<0.0001.

**Supplemental Figure 4 (related to Figure 5).**
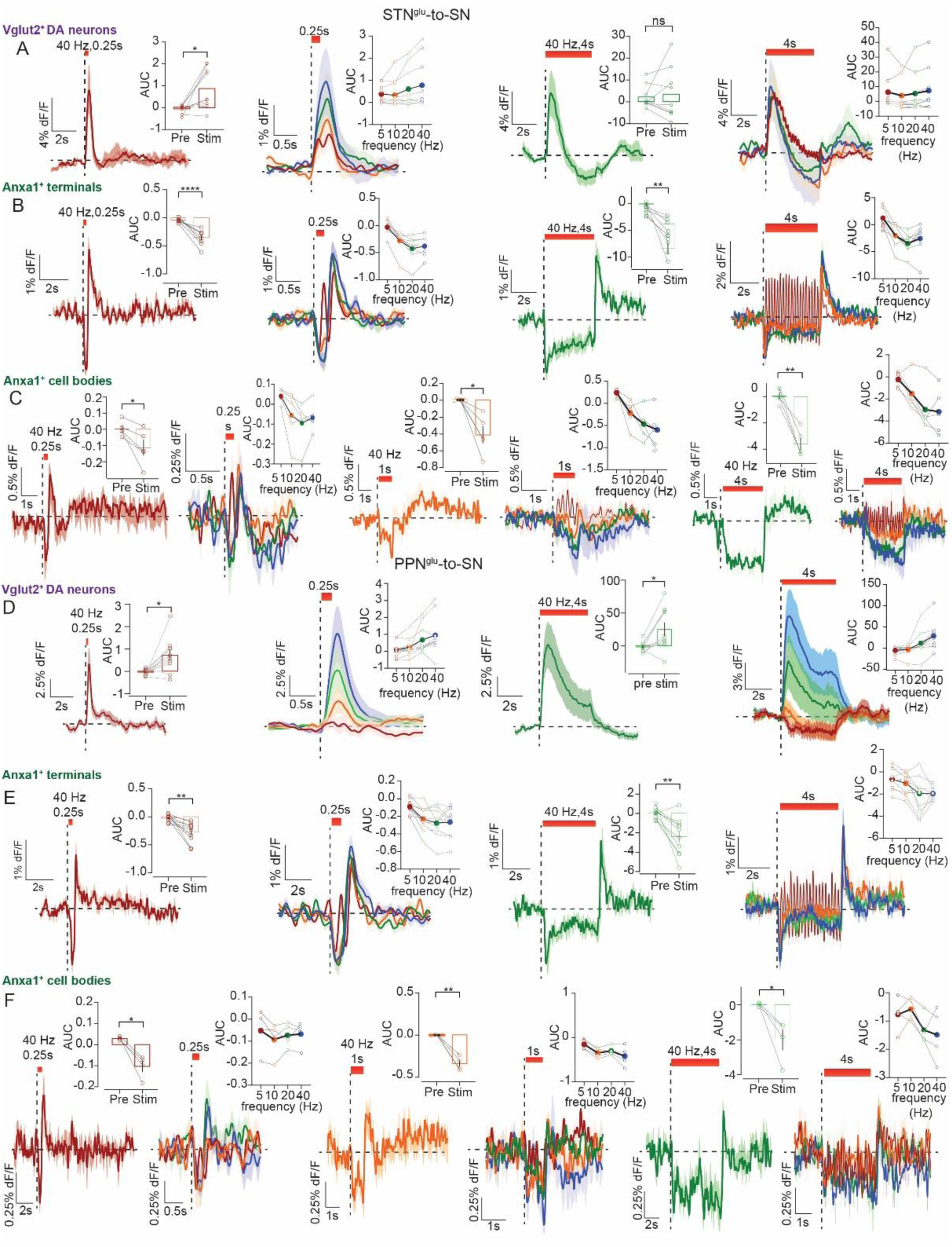
Comparing DA subtype responses to STN and PPN stimulation across additional parameters. **A)** Stimulation of STN inputs to SN with GCaMP8f recordings from Vglut2+ DA neurons (n=10). **B)** Stimulation of STN inputs to SN plus GCaMP8f recordings from Anxa1+ SN terminals in DLS (n=8). **C)** Stimulation of STN inputs to SN plus GCaMP8f recordings from Anxa1+ soma in SNc (n=5). **D)** Stimulation of PPN inputs to SN with GCaMP8f recordings from Vglut2+ DA neurons (n=10). **E)** Stimulation of PPN inputs to SN plus GCaMP8f recordings from Anxa1+ SN terminals in DLS (n=10). **F)** Stimulation of PPN inputs to SN plus GCaMP8f recordings from Anxa1+ soma in SNc (n=4). Paired t-test or Wilcoxon, *p<0.05, **p<0.01, ****p<0.0001

**Supplemental Table 1.**
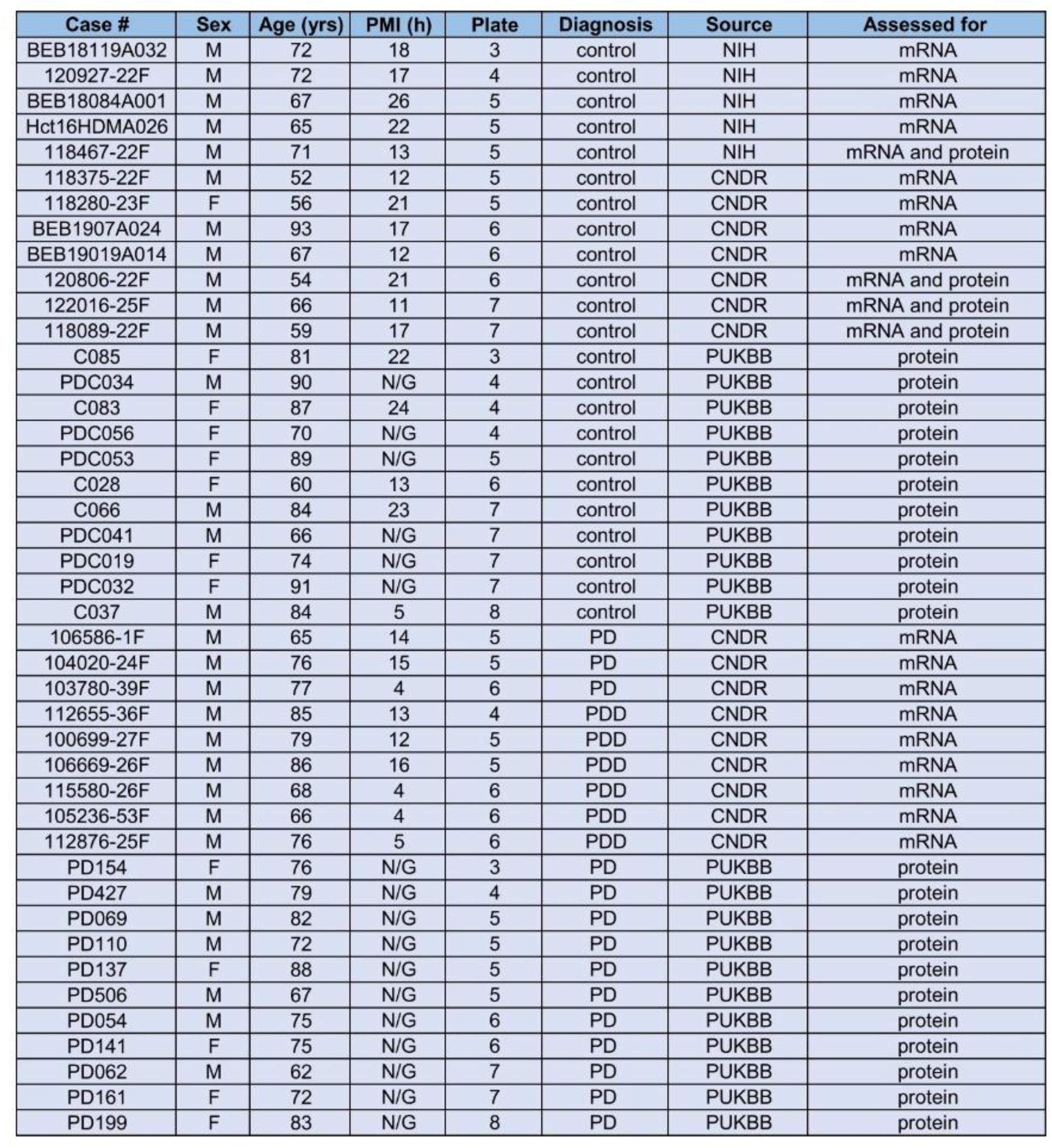
Demographic and other data for cases assessed for mRNA, protein, or both. Level is based on the human brain atlas published by Coulomb et al. 2021 Front Neuroanat. Abbreviations: PD, Parkinson’s disease; PDD, Parkinson’s disease with dementia; NIH, National Institutes of Health NeuroBioBank; CDNR, Center for Neurodegenerative Disease Research at the University of Pennsylvania; PUKBB Parkinson’s United Kingdom Brain Bank; N/G, not given.

**Supplemental Table 2.**
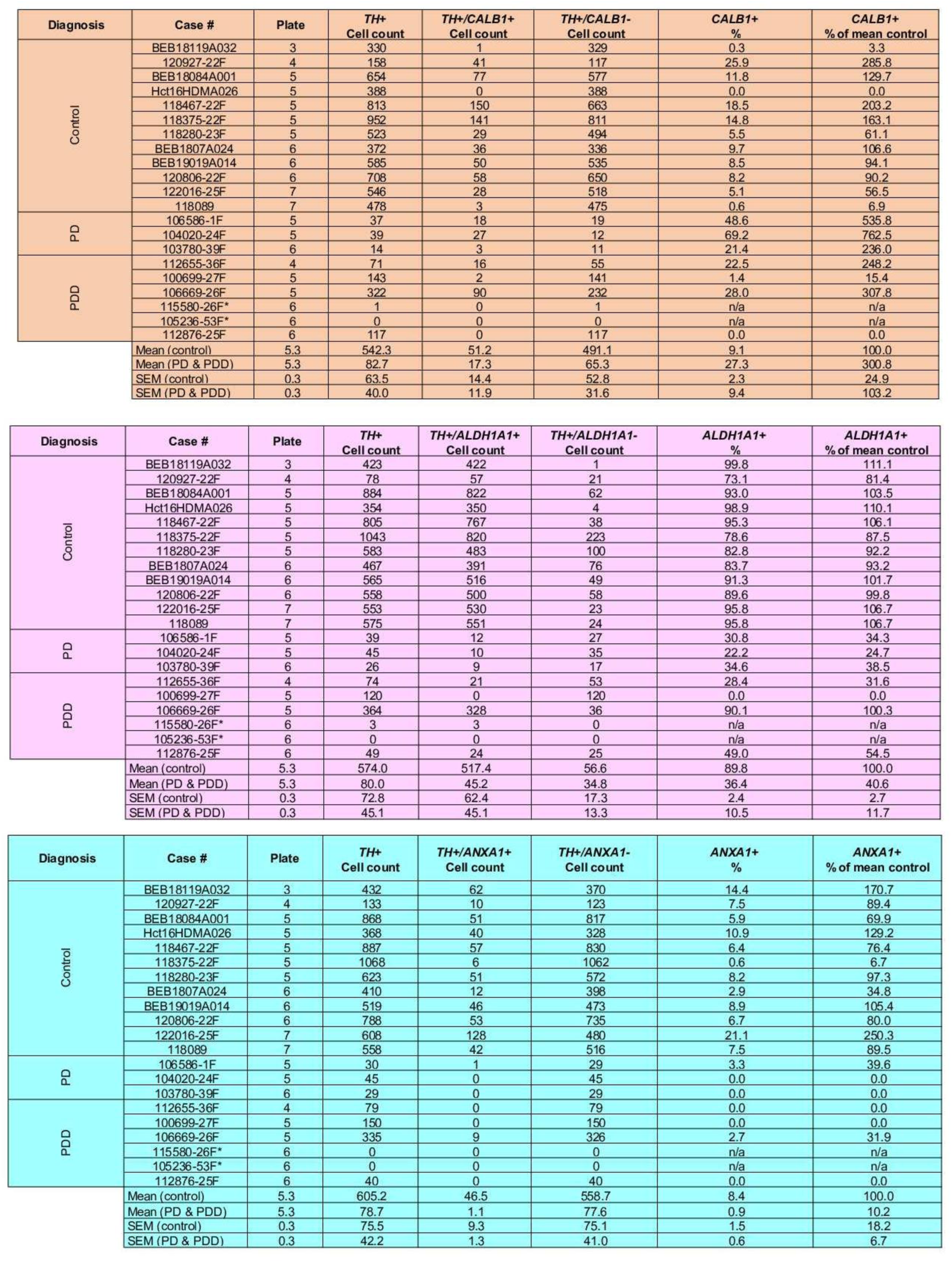
*TH*, *CALB1*, *ALDH1A1*, and *ANXA1* neuron counts in SNc of control and age-matched PD cases with RNAscope labeling. *Cases excluded from analyses because the total number of *TH*+ cells counted was too low.

**Supplemental Table 3.** TH and ALDH1A1 in SNc of control and age-matched PD cases with immunofluorescence labeling.

| Diagnosis | Case # | Plate | TH+<br>Cell count | TH+/ALDH1A1+<br>Cell count | TH+/ALDH1A1-<br>Cell count | ALDH1A1+<br>% | ALDH1A1+<br>% of mean control |
| --- | --- | --- | --- | --- | --- | --- | --- |
| Control | C085 | 3 | 132 | 120 | 12 | 90.9 | 110.3 |
|  | PDC034 | 4 | 282 | 223 | 59 | 79.1 | 96.0 |
|  | C083* | 4 | 302 | 263 | 39 | 87.1 | 105.7 |
|  | PDC056 | 4 | 332 | 269 | 63 | 81.0 | 98.3 |
|  | PDC053 | 5 | 156 | 130 | 26 | 83.3 | 101.1 |
|  | C028 | 6 | 362 | 277 | 85 | 76.5 | 92.9 |
|  | C066 | 7 | 301 | 190 | 111 | 63.1 | 76.6 |
|  | PDC041 | 7 | 716 | 600 | 116 | 83.8 | 101.7 |
|  | PDC019 | 7 | 668 | 558 | 110 | 83.5 | 101.4 |
|  | PDC032 | 7 | 322 | 269 | 53 | 83.5 | 101.4 |
|  | C037 | 8 | 389 | 331 | 58 | 85.1 | 103.3 |
|  | 118375 | 5 | 339 | 232 | 107 | 68.4 | 83.1 |
|  | 118089 | 7 | 276 | 274 | 2 | 99.3 | 120.5 |
|  | 118467 | 5 | 354 | 290 | 64 | 81.9 | 99.4 |
|  | 122016 | 7 | 273 | 243 | 30 | 89.0 | 108.0 |
| PD | PD154 | 3 | 52 | 8 | 44 | 15.4 | 18.7 |
|  | PD427 | 4 | 51 | 35 | 16 | 68.6 | 83.3 |
|  | PD069 | 5 | 57 | 27 | 30 | 47.4 | 57.5 |
|  | PD110 | 5 | 38 | 14 | 24 | 36.8 | 44.7 |
|  | PD137 | 5 | 33 | 14 | 19 | 42.4 | 51.5 |
|  | PD506 | 5 | 72 | 40 | 32 | 55.6 | 67.4 |
|  | PD054 | 6 | 203 | 118 | 85 | 58.1 | 70.5 |
|  | PD141 | 6 | 65 | 27 | 38 | 41.5 | 50.4 |
|  | PD062 | 7 | 38 | 12 | 26 | 31.6 | 38.3 |
|  | PD161 | 7 | 100 | 44 | 56 | 44.0 | 53.4 |
|  | PD199 | 8 | 21 | 16 | 5 | 76.2 | 92.5 |
|  | Mean (control) | 5.7 | 350.2 | 284.6 | 62.3 | 82.4 | 100.0 |
|  | Mean (PD & PDD) | 6.0 | 66.4 | 32.3 | 34.1 | 47.1 | 57.1 |
|  | SEM (control) | 0.4 | 57.0 | 49.9 | 11.4 | 6.1 | 7.4 |
|  | SEM (PD & PDD) | 0.4 | 23.2 | 14.0 | 9.5 | 5.1 | 6.2 |

